# Injury-induced inflammatory signaling and hematopoiesis in *Drosophila*

**DOI:** 10.1101/2021.10.13.464248

**Authors:** Cory J. Evans, Ting Liu, Juliet R. Girard, Utpal Banerjee

## Abstract

Inflammatory response in *Drosophila* to sterile (axenic) injury in embryos and adults has received some attention in recent years, and most concentrate on the events at the injury site. Here we focus on the effect sterile injury has on the hematopoietic organ, the lymph gland, and the circulating blood cells in the larva, the developmental stage at which major events of hematopoiesis are evident. In mammals, injury activates Toll-like receptor (TLR)/NFκB signaling in macrophages, which then express and secrete secondary, pro-inflammatory cytokines. In *Drosophila* larvae, distal puncture injury of the body wall epidermis causes a rapid activation of Toll and Jun kinase (JNK) signaling throughout the hematopoietic system and the differentiation of a unique blood cell type, the lamellocyte. Furthermore, we find that Toll and JNK signaling are coupled in their activation. Secondary to this Toll/JNK response, a cytokine, Upd3, is induced as a Toll pathway transcriptional target, which then promotes JAK/STAT signaling within the blood cells. Toll and JAK/STAT signaling are required for the emergence of the injury-induced lamellocytes. This is akin to the derivation of specialized macrophages in mammalian systems. Upstream, at the injury site, a Duox- and peroxide-dependent signal causes the activation of the proteases Grass and SPE needed for the activation of the Toll-ligand Spz, but microbial sensors or the proteases most closely associated with them during septic injury are not involved in the axenic inflammatory response.

## Introduction

Breach of the *Drosophila* larval epidermis by septic injury elicits innate immune responses that have been studied in great detail in adult stages where it is systemically mediated, almost entirely, by the fat body *via* the Toll and Imd pathways (reviewed in Anderson 2000; Lemaitre and Hoffman 2007; Valanne *et al.* 2011; Buchon *et al.* 2014). Such immune responses are less well-studied in the larval stages (reviewed in Buchon *et al.* 2014). However, an extensive literature describes parasitization by wasps, which is a uniquely larval response that involves several blood cell-types, including the newly induced lamellocytes. Rarely seen in healthy larvae, these are relatively large, flattened cells that function in barrier formation around the parasitoid wasp eggs (reviewed in Letourneau *et al.* 2016). The mature blood cell types that dominate the larva during homeostasis are the macrophage-like plasmatocytes (95%) that secrete antimicrobial peptides, remove cellular debris, and remodel tissue and their extracellular matrix (ECM). The third cell type is crystal cells that provide enzymes important for blood clotting and melanization and aid in injury resolution and antimicrobial responses. As in all invertebrates, *Drosophila* blood cells are functionally similar to those from the vertebrate myeloid lineage. Early genetic studies have shown that constitutive activation of Toll signaling causes blood cell proliferation, precocious lamellocyte differentiation, and melanization responses in larvae (Govind 1996; Qiu *et al.* 1998).

Hemocytes (blood cells) that arise from the head-mesoderm during embryonic development, constitute a circulating and a sessile population of cells underneath the larval cuticle. Additionally, the major site of hematopoiesis in the larva is the specialized blood-forming organ, the lymph gland, which also originates in the embryo, but matures during the larval stages, serving as a hematopoietic reservoir for the pupal and adult stages (reviewed in Banerjee *et al.* 2019). At the onset of metamorphosis, the lymph gland disintegrates, releasing blood cells into circulation. Both larval circulating cells and lymph gland-derived cells comprise the adult fly hemocyte population (Holz *et al.* 2003).

This manuscript probes the effects of axenic or sterile injury to the larvae and the cascade of events that follow, affecting the lymph gland and the circulating hemocytes. This is not the first study on sterile injury in *Drosophila*, but it is designed to probe a different aspect of the larger problem and thereby fulfills an important gap in our knowledge. First, other studies involve embryos or adults (Wood *et al.* 2002; Ramet *et al.* 2002; Galko and Krasnow 2004; Mace *et al.* 2005; Stramer *et al.* 2005; Chakrabarti *et al.* 2020), and not the larval stages where the majority of active hematopoiesis that will populate the pupal and adult stages occurs. Second, the emphasis in other studies is on events at the injury site and in its wound healing (Wood *et al.* 2002; Ramet *et al.* 2002; Galko and Krasnow 2004; Mace *et al.* 2005; Stramer *et al.* 2005; Pastor-Pareja *et al.* 2008; Amcheslavsky *et al.* 2009; Jiang and Edgar 2009; Babcock *et al.* 2009; Chatterjee and Ip 2009; Nam *et al.* 2012; Chakrabarti *et al.* 2016; Chakrabarti *et al.* 2020). Here, we concentrate on the molecular events that mediate the long-distance effect of remote sterile injury on the blood cells and the developing lymph gland, rather than how the macrophages and crystal cells help repair the injury site. The focus of previous studies in the larva, on melanization, lamellocyte formation, and transcriptomic analysis (Goto *et al.* 2003; Bidla *et al.* 2007, Rizki and Rizki, 1992; Márkus *et al.* 2005; Ramond *et al.* 2020), was not on the lymph gland, or on the detailed mechanistic interaction between pathways that result in the hematopoietic response. The results presented here are consistent (but not redundant) with those seen in gut stem cells and/or inferred from survival, when adult flies are injured (Chakrabarti *et al.* 2016; Chakrabarti *et al.* 2020). Altogether, it seems clear that although *Drosophila* has an open circulatory system, the responses to sterile injury are highly conserved and resemble those for vertebrate inflammatory pathways (reviewed in Niethammer 2016).

In mammals, tissue-resident macrophages and fast-responding neutrophils sense inducers of injury and inflammatory responses. They mediate local inflammation through the secretion of secondary proinflammatory signals (Medzhitov 2010; Liu *et al.* 2017). In the case of microbial infection, inductive signals are referred to as pathogen-associated molecular patterns (PAMPs) whereas, for injury, they are the damage-associated molecular patterns (DAMPs) (Kono and Rock 2008; Newton and Dixit 2012). Blood cells utilize so-called pattern recognition receptors (PRRs) to sense both PAMPs and DAMPs. In mammals, a major class of PRRs is the Toll-like receptor (TLR) family, thus named for its shared homology with *Drosophila* Toll (Lemaitre *et al.* 1996).

In *Drosophila*, PAMPs (such as β-glucan and peptidoglycan) are sensed by soluble PRRs (such as GNBP1 and GNBP3) in the hemolymph, and they initiate a proteolytic signaling cascade (including Spaetzle processing enzyme and Grass) that eventually activates a cytokine-like protein ligand, Spaetzle (Spz). Activated Spz binds Toll causing a translocation of the NFκB-like proteins Dorsal and/or Dorsal-related immunity factor (Dif) to the nucleus (Ip *et al.* 1993; Wu and Anderson 1998). The Dorsal/Dif transcription factors upregulate the expression of many downstream genes including antimicrobial peptides (AMPs). While much is understood about the humoral function of Toll in the fat body, relatively little is known about Toll signaling in *Drosophila* blood cells. We report here, the order and timing of events that link injury to Toll activation in blood cells and its downstream consequences on secondary signals and hematopoiesis.

## Results

### Sterile injury activates Toll signaling in the hematopoietic system

Injury caused by posterio-lateral body wall puncture of wandering third-instar larvae causes robust melanization of the wound area by three hours post injury (3 hpi, Figure 1A). To explore Toll pathway signaling, we first assessed the activation of *Drosomycin* (*Drs-GFP*), a classic reporter for Toll signaling in the fat body in response to infection (Lemaitre *et al.* 1996; Ferrandon *et al.* 1998) and found that distal epidermal injury with a sterilized needle is sufficient to activate *Drs-GFP* in both circulating and lymph gland hemocytes by 3 hpi (Figure 1B-E). The injury does not puncture the gut or the lymph gland. We also employed a multigenerational regimen of larval axenic growth condition and tested the larval extract for lack of bacterial growth prior to injury with the sterile needle (see methods; Brummel *et al.* 2004). Furthermore, we used a second synthetic Toll pathway reporter called *D4-lacZ*, containing four copies of known Dorsal binding sites from the *zen* enhancer (Pan and Courey 1992). This reporter gives results similar to those seen with *Drs-GFP*, which also contains Dl binding sites (Sup. Figure S1A-B). Moreover, we find that by 3 hpi, *D4-lacZ* is robustly upregulated in circulating and lymph gland cells of the larvae treated under axenic injury conditions (Figure 1F-I). Consistent with these sterile puncture results, we also find that pinch wounding of the epidermis, which abrades the epidermal cell layer and its underlying ECM without disrupting the outer cuticle (Galko and Krasnow 2004), also causes a similar, though less robust, up-regulation of *D4-lacZ* expression (not shown). Taken together, we conclude that the Toll/Dl pathway is an inflammatory response that is activated even without the mediation of a pathogenic infection.

**Figure 1.**
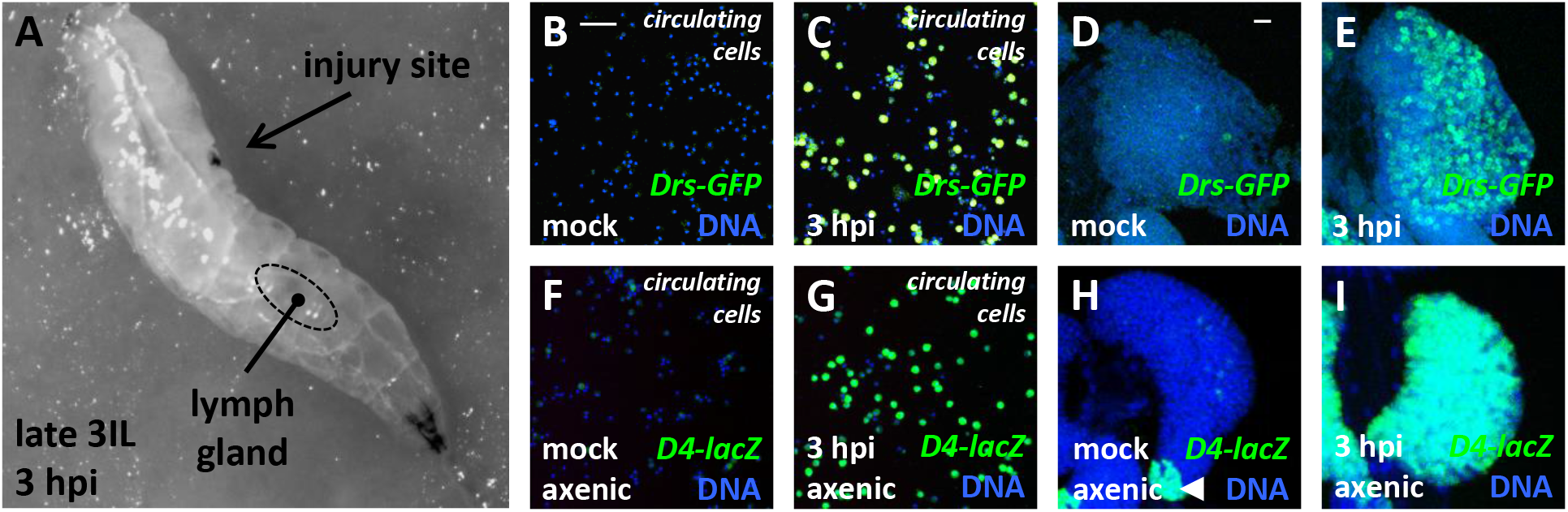
Activation of NFκB-like signaling in the blood system in response to injury. **(A)** Third instar larva showing the location of a melanized puncture wound (arrow) relative to the location of the lymph gland (dotted circle). **(B-I)** Expression of *Drs-GFP* (**B-E**; green) and *D4-lacZ* (**F-I;** green), both of which are reporters for Toll signaling. DNA (blue) staining marks all cells. **(B-C**, **F-G)** Circulating cells. Scale bars 50 μm. **(D-E, H-I)** lymph gland (primary lobes). Scale bars 20 μm. **(C, E)** Injury with a sterile needle; **(G, I)** Axenically reared larvae injured with a sterile needle. Expression of both reporters is low under mock (uninjured) conditions **(B, D, F, H)** but are both induced 3 hours post injury (3 hpi) in the circulating cells as well as in the lymph gland **(C, E, G, I)**. PSC cells constitutively express *D4-lacZ* (arrowhead in **H**). In **Sup. Figure 1K-O**, we show that Dorsal is not expressed in PSC cells, but Dif is, and the expression of the reporter is lost in a *Dif* mutant even though the PSC remains intact.

The expression patterns of *Drs-GFP* and *D4-lacZ* also accurately reflect the subcellular localization of Dorsal. In uninjured animals, an antibody raised against the Dorsal protein shows robust cytoplasmic localization in both circulating and lymph gland cells (Sup. Figure 1C, E). Interestingly, in the lymph gland, this cytoplasmic localization is largely seen in the progenitors of the medullary zone (Sup. Figure 1E-F). Following sterile injury, Dorsal protein is now found in the nucleus of a great majority of these cells by 3 hpi (Sup. Figure 1D, F). The activation of *D4-lacZ* in response to injury is completely obliterated in a genetic background that is deficient for both Dorsal and Dif gene products (Sup. Figure 1G-H), or upon the loss of the intracellular Toll signaling adapter protein Myd88 (Sup. Figure 1I-J). Thus the Toll pathway and its downstream NFκB-like protein(s) control the sterile inflammatory response of the larval hemocytes. Given that Toll is autonomously required, the wound site must communicate with the blood cells by a systemic mechanism.

### Injury-induced Toll signaling in the blood requires Spaetzle

Spz is the only known ligand in *Drosophila* for the Toll receptor (Lemaitre *et al.* 1996; Weber *et al.* 2003; Parthier *et al.* 2014). We find that injured heterozygous control larvae (*spz^rm7^/+*) exhibit normal *D4-lacZ* activation upon injury (Figure 2A), however when the *spz^rm7^* mutant allele is placed over a chromosomal deficiency for the *spz* locus (*spz^rm7^/Df spz*), the activation of *D4-lacZ* in response to sterile injury is completely suppressed (Figure 2B). Additionally, qRT-PCR analysis demonstrates that injury induces the expression of the endogenous *Drosomycin* (*Drs*) gene, and loss of *spz* function completely suppresses this response (Sup. Figure 2A). Consistent with these data, injured control larvae (*spz^rm7^/+*) accumulate Dorsal in the nucleus of blood cells in circulation and in the lymph gland by 3 hpi (Sup. Figure 2B, D, F), whereas injured *spz* mutant larvae maintain Dorsal in the cytoplasm of these cells over the same time period (Sup. Figure 2C, E, G). These data establish that injury-induced Toll pathway signaling in blood cells requires the ligand Spz.

**Figure 2.**
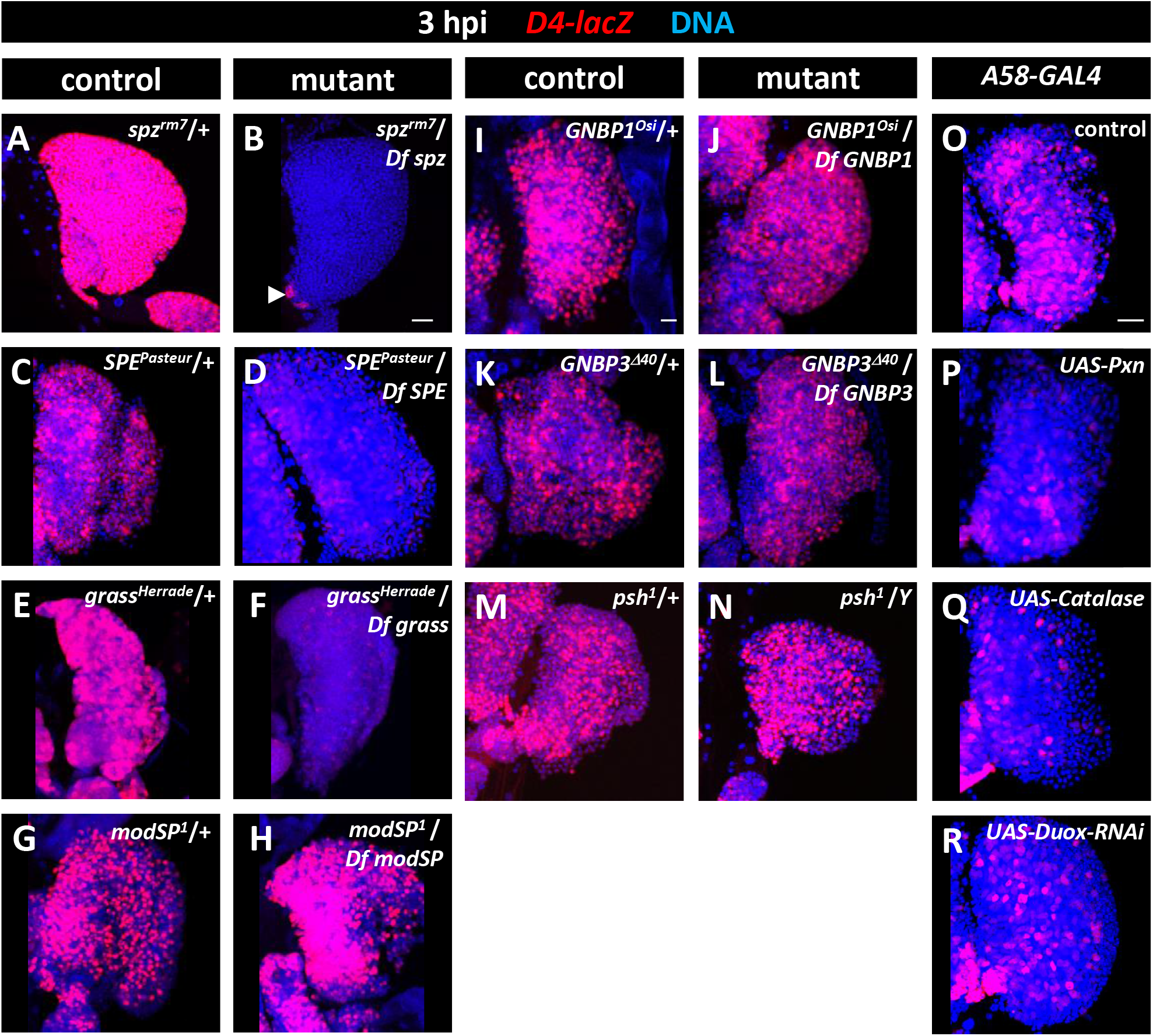
The Toll pathway serine proteases Grass and SPE are required to activate Spz downstream of ROS for injury-induced signaling in blood cells. **(A-R)** *D4-lacZ* (Toll pathway reporter, red) in lymph glands 3 hpi; DNA (blue); Scale bars 20 μm. **(A-F)** Toll reporter expression is robustly activated in the lymph gland by 3 hpi in heterozygous control animals **(A, C, E)**, but is completely suppressed in *spz, SPE*, or *grass* mutant lymph glands **(B, D, F)**. Note that *D4-lacZ* expression (arrowhead in **B**) is retained in PSC cells in this mutant background (compare with **Figure 1H**). **(G-N)** Mutants in proteases *modSP* **(H)** and *psh* **(N),** or PRRs *such as GNBP1* **(J)** and *GNBP3* **(L)** that function upstream of Grass do not affect injury-induced *D4-lacZ* expression compared to heterozygous controls **(G, I, K, M)**. **(O-P)** Targeted expression in epidermal cells (*A58-GAL4*) of scavengers of ROS, Pxn **(P)** or human Catalase **(Q)** suppresses injury-induced *D4-lacZ* expression in the lymph gland relative to control **(O)**. **(R)** Specific knockdown of Duox function (*A58-GAL4, UAS-Duox^RNAi^*) also suppresses *D4-lacZ* activation in response to injury in the lymph gland relative to control **(O)**. This establishes that Duox mediates ROS-dependent Toll signaling in blood cells.

### The Toll-mediated injury response in blood requires the proteases SPE and Grass

The active Spz ligand that mediates Toll receptor signaling is produced through proteolytic processing of a Spz pro-protein (Pro-Spz; Schneider *et al.* 1994). During immune challenge in larvae and adults, Pro-Spz is converted into active Spz by the serine protease Spaetzle Processing Enzyme (SPE; Valanne *et al.* 2011). Similar to what is seen for Spz, loss of SPE activity (*SPE^Pasteur^/Df SPE*) also strongly suppresses *D4-lacZ* activation in blood cells in response to injury (Figure 2C-D). In the context of infection, the sensing of Gram-positive bacteria leads to the sequential activation of the serine proteases ModSP, Grass, and then SPE (Jang *et al.* 2006; El Chamy *et al.* 2008; Buchon *et al.* 2009a). Whereas, in the context of injury, while loss of Grass (*grass^Herrade^/Df grass*) strongly suppresses *D4-lacZ* expression (Figure 2E-F), loss of ModSP function (*modSP^1^/Df modSP*) does not (Figure 2G-H). Thus, sterile injury-induced activation of Spz and downstream Toll signaling in the blood system requires the two proteases SPE and Grass, that are immediately upstream of Spz, but not the further upstream protease ModSP, which functions in the context of infection.

The activation of Spz/Toll in D/V patterning of the embryo requires an entirely different set of proteases. These are Easter (Ea), Snake (Snk), Gastrulation defective (Gd), and Nudel (Ndl). Using the D4-lacZ sterile injury assay, we are able to rule out a function of this entire cascade since mutations in *ea (ea^4^/Df ea), snk (snk^2^/Df snk), gd (gd^1^/Df gd*), or *ndl* (*ndl^10^/Df ndl*) exhibit no suppression of sterile injury-induced *D4-lacZ* expression (Sup. Figure 2H-O). This indicates that the embryonic, developmental Toll signaling cohort is not involved in inflammatory response.

### Sterile injury response is independent of microbe sensors but requires injury-site production of hydrogen peroxide

The pattern recognition receptor (PRR) GNPB1 is essential for sensing Lysine-type peptidoglycan (Lys-PGN) produced by Gram-positive bacteria and assists in the downstream activation of ModSP and Grass (Gobert *et al.* 2003; El Chamy *et al.* 2008; Buchon *et al.* 2009a). Sterile injury does not involve pathogens and accordingly, we detect no role for GNBP1 in this process (Figure 2I-J). Additional PRRs that sense pathogens such as GNBP3 (Gottar *et al.* 2006) and Persephone (Psh; Ligoxygakis *et al.* 2002) also do not affect post-injury induction of *D4-lacZ* (Figure 2K-N). Thus, while Grass, SPE, and Spz are common to both injury- and infection-induced Toll signaling, the mechanism of injury sensation leading to Grass activation uses different, possibly unique, components.

In both zebrafish and *Drosophila* embryos and adults, hydrogen peroxide produced at injury sites serves as a chemoattractant for migrating blood cells (Niethammer *et al.* 2009; Moreira *et al.* 2010; Chakrabarti *et al.* 2020). To determine if injury-site-derived hydrogen peroxide also plays a role in Toll pathway activation in the blood, we enhanced peroxide scavenging in the epidermis with the epidermis-restricted expression of *Drosophila* Peroxidasin (*A58-GAL4 UAS-Pxn;* Figure 2P) or the human Catalase enzyme (*A58-GAL4 UAS-Cat;* Figure 2Q), and we found that the injury-induced Toll signaling (assessed by *D4-lacZ* expression) in the distant blood cells is markedly reduced (Figure 2O-Q).

In the *Drosophila* embryo, immediately following injury, a flash of Ca^2+^ propagates outward from the wound site and activates the Dual oxidase (Duox) enzyme that generates hydrogen peroxide (Ha *et al.* 2009; Moreira *et al.* 2010; Juarez *et al.* 2011; Razzell *et al.* 2013. Similar phenomena have also been reported in vertebrates (Niethammer *et al.* 2009; Yoo *et al.* 2011). In the context of larval epidermal injury, we find that knockdown of Duox activity in the epidermis (*A58-GAL4 UAS-Duox RNAi*) suppresses the injury-induced Toll signaling in blood cells (Figure 2O, R), similar to that seen when ROS is scavenged in the epidermis (Figure 2O-Q). Collectively, these data indicate that injury site production of hydrogen peroxide by Duox is essential for the systemic activation of Toll signaling in the *Drosophila* blood.

### Epidermal injury causes Toll pathway-dependent hematopoietic differentiation of lamellocytes

Hematopoietic differentiation of lamellocytes is closely associated with the cellular immune response to parasitoid wasp infestation of larvae (Rizki and Rizki 1992; Carton and Nappi 1997; Sorrentino *et al.* 2002; Márkus *et al.* 2005). Importantly, wasp oviposition also causes a puncture injury to the larval body wall including the overlying cuticle, epidermis, and the underlying ECM layer as it delivers the wasp embryo and venom into the larval hemocoel. Subsequently, lamellocytes differentiate and, along with plasmatocytes and crystal cells, encapsulate the parasitic embryo in an effort to destroy it (reviewed in Letourneau *et al.* 2016). With our experimental approach, we likewise find that larvae injured with a sterile needle exhibit an increase in lamellocyte number in lymph glands and in circulation by 24 hpi, relative to uninjured control larvae (Figure 3A-E). Additionally, we find that injury-induced lamellocyte formation is strongly reduced both in lymph glands and in circulation when *spz* function is lost (24 hpi, Figure 3E). These results indicate that the rapid Toll pathway signaling observable throughout the hematopoietic system by three hours post-injury is consolidated over time with other molecular events such as lamellocyte differentiation that takes place almost a day after the injury. We also conclude that while the wasp egg and its associated venom might have some role in this process, the trigger for lamellocyte induction is the breach of epidermis, and not the presence of the parasite.

**Figure 3.**
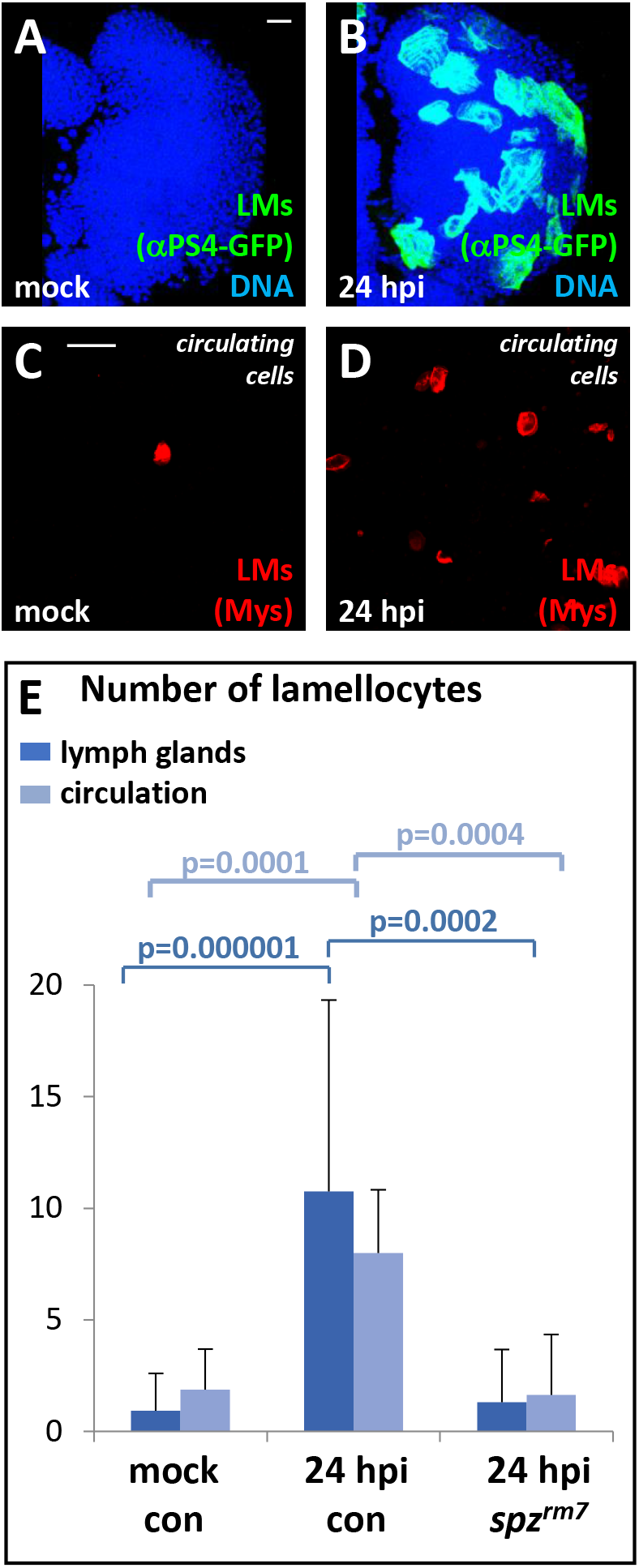
Injury-induced lamellocyte differentiation requires Spz. Lamellocytes (LMs) differentiate in lymph glands by 24 hpi **(A-B)**, identified by the specific expression of the αPS4-GFP reporter (green). DNA (blue); Scale bars 20 μm. Lamellocyes also differentiate among circulating cells **(C-D)**, identified by their high expression of Mys (red), also by 24 hpi. Scale bars: 100 μm. Mutation in *spz* (*spz^rm7^/spz^rm7^*) completely blocks injury-induced lamellocyte differentiation, with lamellocyte counts similar to that in control **(E)**.

### Epidermal injury activates Jun kinase signaling in the blood system in a Toll pathway-dependent manner

Classic genetic analysis has shown that lamellocyte differentiation relies on activation of the JNK pathway (Zettervall *et al.* 2004; Tokusumi *et al.* 2009), and we asked whether this is also true in the context of injury. *puc-lacZ* is a canonical JNK signaling reporter (Martín-Blanco *et al.* 1998), and a mild, but discernible, increase in its expression is apparent in lymph gland cells by 3 hpi (Figure 4A-B). A second widely studied JNK target encodes *Matrix metalloproteinase 1* (*Mmp1*), a secreted protease with developmental and ECM remodeling roles (Uhlirova and Bohmann 2006; Srivastava *et al.* 2007; Page-McCaw *et al.* 2007; Stevens and Page-McCaw 2012). In contrast to the weak *puc-lacZ* induction, Mmp1 protein is robustly up-regulated at the early, 3 hpi time point (Figure 4C-D). Epidermal injury also upregulates both the endogenous *Mmp1* mRNA in circulating cells outside of the lymph gland (Figure 4E) as well as *Mmp1-lacZ* reporter expression in lymph glands by 3 hpi (Figure 4F, H). *Drosophila* Jun kinase (Basket; Bsk), is activated by the corresponding JNK kinase (JNKK) protein encoded by *hemipterous* (*hep*) (Glise *et al.* 1995), and in the genetic background of a null *hep^r75^* mutant or targeted expression of a dominant-negative Bsk allele (*Hml^Δ^-GAL4 UAS-bsk^DN^*), expression of *Mmp1-lacZ* is no longer up-regulated post-injury (Figure 4F-I). Thus, injury causes Dorsal nuclear localization and JNK activation as early as 3 hpi even though resulting lamellocytes take up to 24h to form.

**Figure 4.**
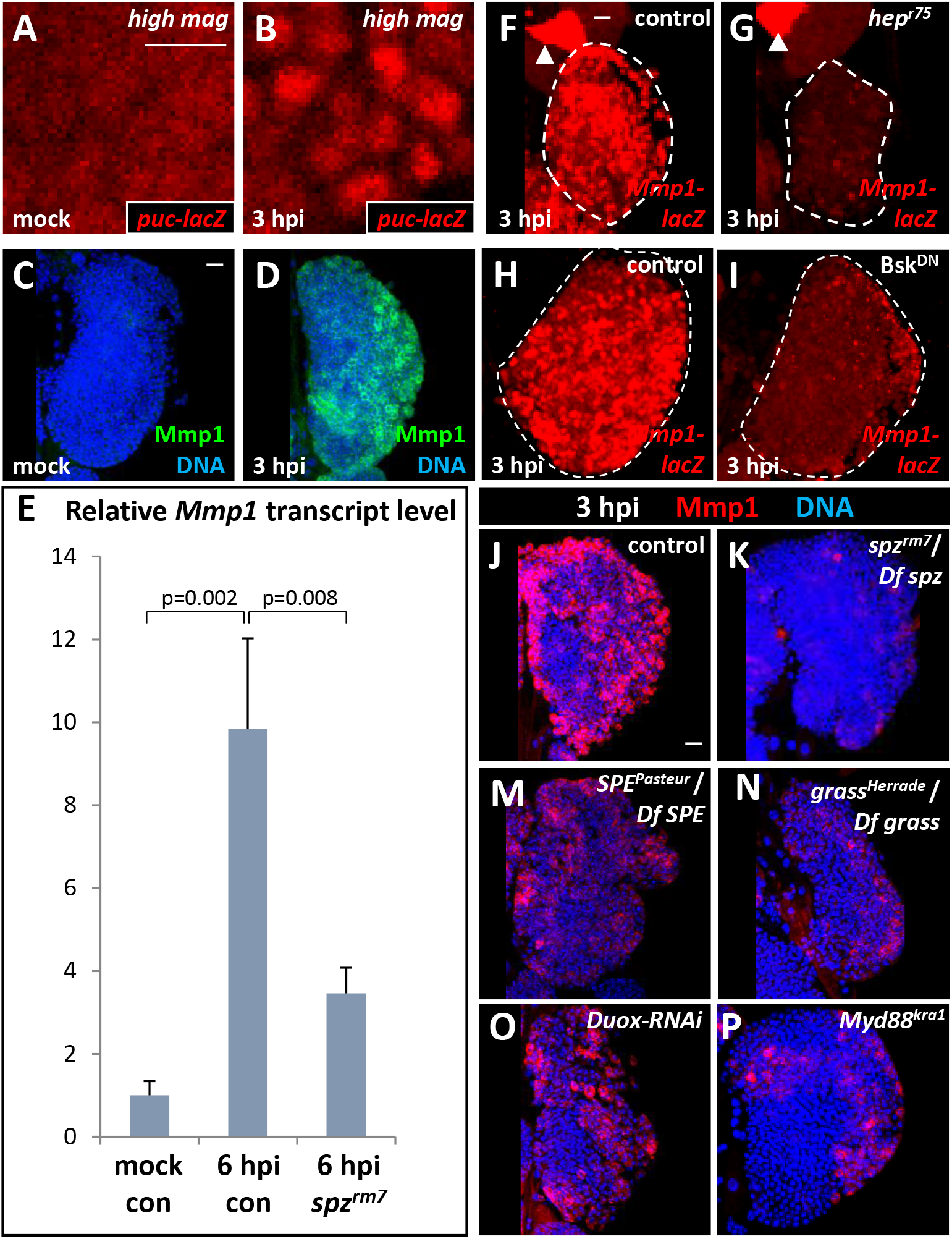
Injury rapidly activates JNK signaling in the lymph gland in a Toll- and epidermal Duox-dependent manner. **(A-B)** Compared with control **(A),** the JNK signaling reporter *puc-lacZ* (red) is upregulated upon sterile injury at 3 hpi **(B)**. Scale bars 20 μm. **(C-D)** The JNK pathway target protein Mmp1 is strongly induced by 3 hpi **(D)** compared with uninjured control **(C)**. DNA (blue); Scale bars 20 μm. **(E)** qRT-PCR results showing that *Mmp1* mRNA transcript levels are robustly induced in circulating cells by 6 hpi, but are much lower in *spz* mutants (n=3). **(F-I)** Injury-induced upregulation of *Mmp1-lacZ* (red) **(F, H)** is suppressed in a *hep^r75^* mutant background **(G)** or by dominant-negative Bsk (*Hml^Δ^-GAL4 UAS-Bsk^DN^;* **I**), both indicating a suppression of the injury-induced JNK activation. The white arrowheads in panels **(F)** and **(G)** show that control *Mmp1-lacZ* expression in ring glands is not affected by the loss of JNK activation. Scale bars 20 μm. **(J-P)** Induction of Mmp1 protein (red) at 3 hpi. DNA (blue); Scale bars 20 μm. **(J-K)** Mmp1 protein is strongly suppressed by a mutation in *spz.* **(M-P)** As with loss of *spz*, loss of *SPE* **(M)** or Grass **(N)** suppresses injury-induced Mmp1 upregulation. Knockdown of Duox in epidermal cells (*A58-GAL4 UAS-Duox-RNAi;* **O**) also suppresses injury-induced Mmp1 expression as does a mutation in *Myd88 (Myd88^kra1^*, **P**), indicating that Toll pathway-mediated JNK activation in response to injury occurs downstream of the Toll/Myd88 complex.

Given its rapid activation, we explored whether the JNK pathway is dependent on, or directly downstream of, Toll signaling. Indeed, we found that loss of *spz (spz^rm7^/Df spz*) strongly suppresses Mmp1 upregulation in the lymph gland upon injury (Figure 4J-K). Also, qRT-PCR analysis of circulating cells post injury show an approximately 60% reduction of *Mmp1* mRNA expression in *spz* mutants (Figure 4E). In Figure 2C-F, R we show that injury-induced Toll signaling in the blood requires the upstream function of SPE, Grass, and Duox. Consistent with these findings, global loss of *Grass (Grass^Herrade^/Df Grass*) or *SPE (SPE^Pasteur^/Df SPE*), or knockdown of Duox in the larval epidermis (*A58-gal4 UAS-Duox-RNAi*), suppresses injury-induced Mmp1 upregulation in the lymph gland (Figure 4J, M-O). This is true as well, with global loss of function of the Toll adapter protein *Myd88* (Figure 4J, P). Thus, both injury-induced JNK signaling and the expression of its downstream target, Mmp1, in blood cells are Toll signaling-dependent, and are rapidly activated downstream of the Toll/Myd88 complex.

### Toll pathway-dependent expression of cytokine-like genes and JAK/STAT activation

In mammalian macrophage activation, the TLR/NFκB signal increases the production of pro-inflammatory cytokines, such as IL-6, IL-12, and TNFα (Medzhitov 2010; Liu *et al.* 2017). In *Drosophila*, the *unpaired* family of genes (*upd, upd2*, and *upd3*) encode cytokine-like ligands that mediate JAK/STAT signaling (Zeidler and Bausek 2013). In particular, Upd3, a four-helix-bundle ligand with homology to IL-6, has been linked to JAK/STAT signaling in blood cells (Agaisse *et al.* 2003; Pastor-Pareja *et al.* 2008; Oldefest *et al.* 2013). We find that *upd3* expression (*upd3-GAL4 UAS-GFP;* Agaisse *et al.* 2003) is up-regulated by injury throughout the lymph gland and in circulating blood cells by 24 hpi (Figure 5A-F), but not at 3 hpi when Toll and JNK signaling is already apparent. When analyzed by qPCR rather than indirectly through marker expression, *upd3* transcripts exhibit a greater than 20-fold increase at 6 hpi (Figure 5G). The expressions of the related cytokine genes *upd* and *upd2* are also similarly upregulated albeit to a lesser extent (Figure 5G), consistent with the reported co-regulation of these genes in many tissues (Pastor-Pareja *et al.* 2008).

**Figure 5.**
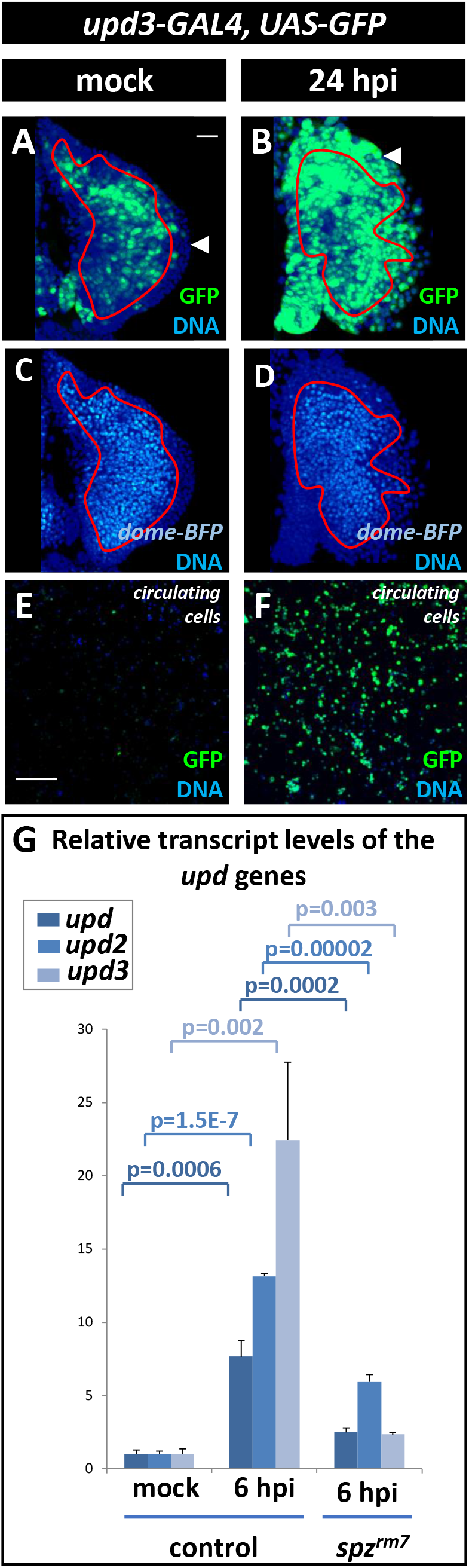
Sterile injury causes a Spz-dependent induction of *upd3* and other cytokine-related genes. **(A-F)** *upd3-GAL4, UAS-GFP* (**A, B, E, F**; green); *dome-BFP* (**C, D**; cyan); DNA (**A-F**; blue). Lymph gland **(A-D)** and circulating hemocytes **(E-F)** are assayed. Medullary zone boundaries (red outlines in **A-D**) are based on *dome-BFP* expression in the same gland (in **C-D**). White arrowheads (in **A-B**) point to the cortical zone. In mock (uninjured) lymph glands **(A, C)**, *upd3-GAL4* is developmentally expressed in the Medullary Zone. Following sterile injury **(B, D)**, by 24 hpi, *upd3-GAL4 UAS-GFP* expression is upregulated in both the Medullary Zone and the Cortical Zone. Circulating hemocytes from uninjured larvae **(E)** do not express *upd3-GAL4*, but by 24 hpi **(F)**, this reporter is strongly upregulated in the circulating hemocytes. Scale bars: 20 μm **(A-D);** 100 μm **(E-F)**. **(G)** Induction of endogenous *upd, upd2*, and *upd3* mRNAs measured by qRT-PCR. Each is upregulated by 6 hpi, with *upd3* showing the strongest (>20-fold) increase in its mRNA. The injury-induced expression of genes encoding these cytokines is lost in homozygous *spz^rm7^* mutants.

Constitutive activation of Toll signaling (*Toll^10B^* mutants, no injury) causes a robust increase in *upd3* reporter expression (*upd3-GAL4 UAS-GFP;* Sup. Figure 3A-B) suggesting that the Toll pathway functions upstream of Upd3. ModEncode ChIP-seq data (Nègre *et al.* 2011) has identified multiple Dorsal binding regions (Sup. Figure 3C) within the *upd3* enhancer region (Agaisse *et al.* 2003). Importantly, upon injury, *spz* null mutants do not up-regulate the expression of *upd3-GAL4* or any of the endogenous *upd* genes (Figure 5G; Sup. Figure 3D-G).

Injury-induced expression of *upd3* suggests that JAK/STAT signaling could play a role in injury response in hemocytes. To this end, we monitored the expression of an *in vivo* reporter that responds to nuclear STAT (*10XSTAT-GFP;* Bach *et al.* 2007), as well as that of the JAK/STAT target genes *myospheroid (mys;* Issigonis *et al.* 2009) and *chronically inappropriate morphogenesis (chinmo;* Flaherty *et al.* 2010). By 24 hpi, the *10XSTAT-GFP* reporter is elevated relative to cells from uninjured control larvae (Figure 6A-B). Likewise, the expression of Mys and Chinmo are upregulated in circulating cells by 24 hpi (Figure 6D-E, G-H). Importantly, qPCR assays show that RNA levels for downstream components *mys* and *chinmo* are upregulated as early as 6 hpi (Sup. Figure 3H-I). The increase in STAT targets is lost when a dominant negative version of the Upd3 receptor, Domeless (Dome) is expressed (Figure 6C, F, I). Blocking Dome function in this manner also strongly reduces injury-induced lamellocyte differentiation (Figure 6J-M). We conclude that upon sterile injury the cytokine Upd3 is transcriptionally induced by the Toll pathways, and *via* the activation of JAK/STAT signaling, it promotes lamellocyte differentiation.

**Figure 6.**
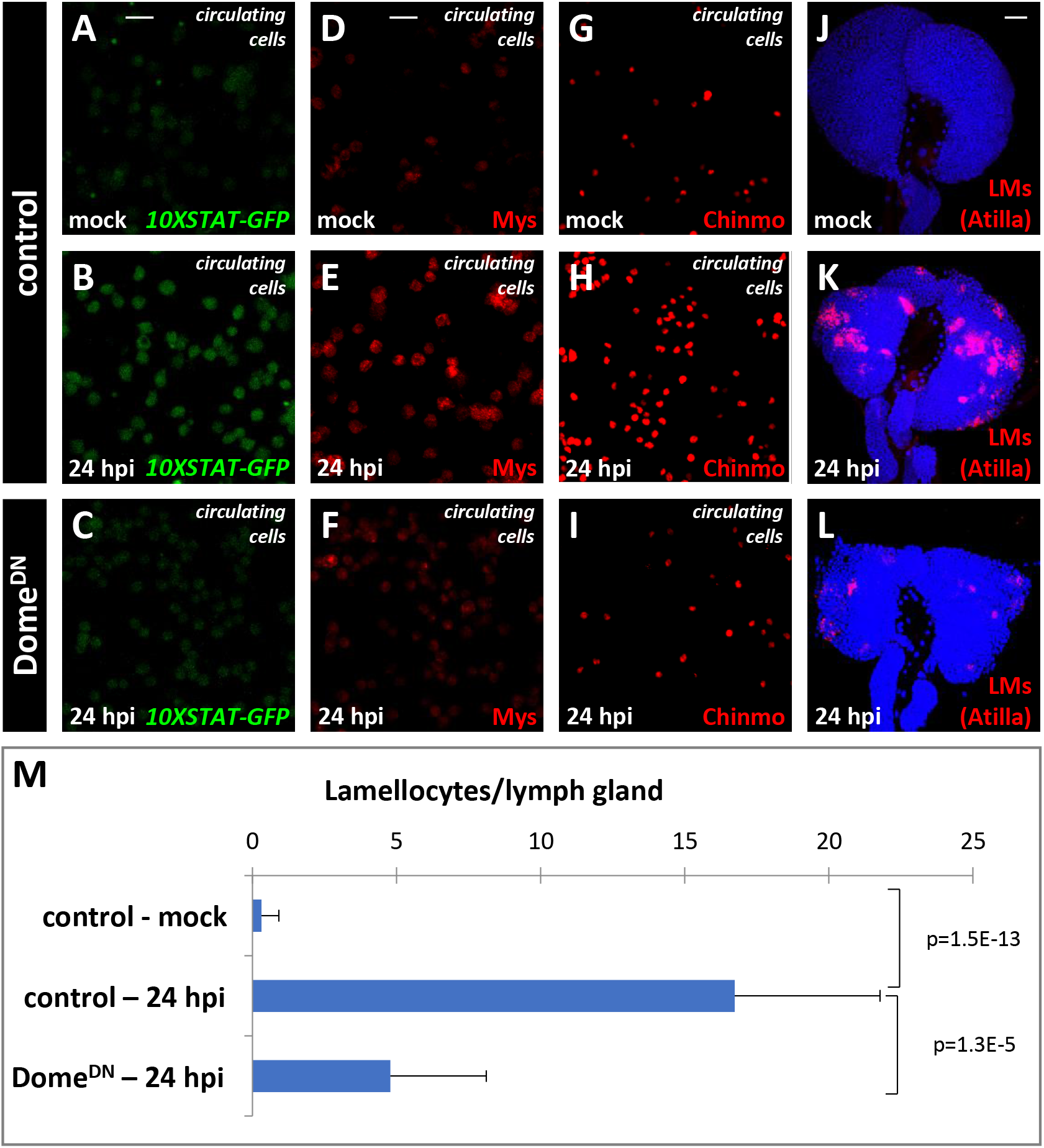
JAK/STAT signaling in blood cells is activated by injury and is required for lamellocyte differentiation. **(A-C)** The JAK/STAT reporter gene *10xSTAT-GFP* (green) is not expressed in circulating hemocytes of uninjured (**A**; mock) larvae. It is induced by 24 hpi **(B)** and suppressed at 24 hpi in larvae expressing a dominant negative form of the receptor Domeless (**C**; *Hml^Δ^-GAL4 UAS-Dome^DN^*). Scale bars: 20 μm. **(D-I)** The same results as in **(A-C)** are also seen for endogenous JAK/STAT target genes *mys* (**D-F**, red) and *chinmo* (**G-I**, red). Scale bars 20 μm. **(J-M)** Blocking the function of the Dome receptor (*Hml^Δ^-GAL4 UAS-Dome^DN^*) also suppresses injury-induced lamellocyte differentiation (Atilla-positive cells, red) in 24hpi lymph glands. Scale bars: 20 μm. Quantitation shown in **M**.

## Discussion

The four classic cardinal signs of tissue inflammation in response to injury: redness (*rubor*), swelling (*tumor*), heat (*calore*), and pain (*dolore*), were described by Celsus in the first century A.D. (Celsus 25AD). In more recent times, molecular and mechanistic analysis has revealed that innate immune response is conserved across evolutionary time (Lemaitre and Hoffmann 2007; Medzhitov 2010; Newton and Dixit 2012). This also seems to be true for the aseptic inflammatory response that is the focus of this study. For example, the major initial phase of Inflammatory response in vertebrates is injury sensation by blood cells and their recruitment to the injury site. In *Drosophila* as well, blood cells are rapidly recruited to sites of injury in both embryos and larvae (Galko and Krasnow 2004; Wood *et al.* 2006; Babcock *et al.* 2008). Cues elicited by physical damage range from small molecules (calcium ions, ATP, ROS, *etc*) to DAMPs derived from cells and ECM, and possibly even protein ligands derived from cells proximal to the wound site (Pastor-Pareja *et al.* 2008; Babcock *et al.* 2009). While the sensation of damage-associated cues and its immediate propagation might vary, it is clear from work in a variety of systems that activation of TLRs and NFκB proteins are important for these processes.

Previous studies of the response to mechanical injury *in Drosophila* have largely focused on deciphering the repair and resolution mechanisms associated with the injury site and, in some cases, how blood cells interact with and facilitate these processes (Wood *et al.* 2002; Mace *et al.* 2005; Babcock *et al.* 2008; Wu *et al.* 2009; Juarez *et al.* 2011; Nam *et al.* 2012; Patterson *et al.* 2013; Chakrabarti *et al.* 2016). Only recently have we started to get some understanding of how wounds change the blood cells themselves (Ramond et al, 2020; Chakrabarti *et al.* 2020). However, prior analyses largely examine circulating cells in the adult and in the embryonic stages although a majority of hematopoietic events occur in larval stages, and the larval cuticle is the most susceptible to injury. The work presented here is therefore focused on two questions: 1. How do *Drosophila* blood cells in the hematopoietic organ (lymph gland) sense and respond to epidermal injury at a distance and 2. What is the sequence of the signaling pathways within the blood cells that explains the changes they experience upon sterile injury.

The Toll pathway and the events that lead up to the activation of its ligand Spz, have been described in significant detail for septic injury and microbial infection, but not, until this current study, for such effects on the hematopoietic system upon sterile injury. Using axenic culture and sterile injury methods, we show that injury alone can lead to activation of Spz by proteolytic cleavage. This pathway utilizes none of the embryonic D/V patterning enzymes that activate Spz. Immunity-related proteases that function close to the Spz-activation cascade (e.g., Grass and SPE) function in the context of sterile injury, but unlike during infection, sterile injury response does not involve the upstream pathogen-sensing components of the response, such as the proteases ModSP and Psh and the sensors GNBP1 and GNBP3. In retrospect, this is to be expected since sterile injury does not involve the sensing of an invading particle. We surmise that since both kinds of injuries involve breach of the epidermis, that septic injury activates two pathways, one from the epidermis and the other originating from the microbe. The two pathways intersect at the level of proximal proteases such as Grass and then follow a common pathway towards Toll activation. The time scales of the two pathways are different with a fast priming injury response followed by a delayed, but much larger response if a microbe is detected.

How epidermal injury activates Grass is not yet fully understood, but our data agree with previous observations in the *Drosophila* embryo and in zebrafish that Duox-mediated hydrogen peroxide (H2O2) produced at the injury site is a critical mediator of this process (Moreira *et al.* 2010; Razzell *et al.* 2013; Chakrabarti *et al.* 2020). A key finding of this work is that this initial event dictates the activation of Toll in distant hemocytes, and their precursors in the hematopoietic organ.

Another important finding is that larval epidermal injury also causes rapid JNK pathway activation within the hematopoietic system, and that this JNK signal is Toll pathway dependent. Mammalian TLR signaling also causes JNK activation (Matsuguchi *et al.* 2003; reviewed in Kawasaki and Kawai 2014), and RNA-Seq results in the context of *Drosophila* injury indicate rapid (by 45 minutes) upregulation of the JNK activation signature (Ramond *et al.* 2020). Nevertheless, previous evidence of such cross-talk in *Drosophila* is relatively scant (Wu *et al.* 2015). In the context of infection in adult flies, Boutros *et al.* (2002) demonstrated the upregulation of JNK pathway target genes, a subset of which were found to be also dependent upon Toll signaling.

Our work indicates that, in the case of injury and specifically in blood cells, Toll signaling activates the JNK pathway in a fairly direct manner. Loss of JNK/JNKK or loss of Myd88 (that complexes with Toll), have the same attenuating effect on JNK target genes such as Mmp1. This result has impressive parallels in mammalian systems, where careful biochemical analysis has identified that a scaffolding protein called JIP3 (JNK interacting protein; Matsuguchi *et al.* 2003) participates in directly associating TLR4 to multiple kinases of the JNK (and MAPK) pathways. Specifically, the authors demonstrate the formation of tripartite complexes of TLR4, JIP3 and JNK. This scaffolding protein is evolutionarily conserved including in *Drosophila* (Bowman *et al.* 2000) and *C. elegans* (Byrd *et al.* 2001). The *Drosophila* protein is called Sunday Driver (SYD). In future studies it will be interesting to see if this model holds up for *Drosophila* sterile injury response by manipulating the *syd* gene. A second as yet unsolved hypothesis in this context is based on the report of Toll/JNK crosstalk in Drosophila (Wu *et al.* 2015). Albeit in a different context, the authors show that the JNK pathway leads (directly or indirectly) to a transcriptional upregulation of Spz. If this were to be true in the context of injury, it will result in a positive feedback loop for Toll activation upon injury. We already know that JNK downstream products like Mmp1 are secreted in response to injury and Toll dependent JNK activation. The possibility of a positive feedback loop will make for an exciting future study.

We have also demonstrated that epidermal injury causes the upregulation of the cytokine-like gene *upd3* in blood cells, and that this occurs along a delayed timeline relative to the other injury-induced markers (6 hpi instead of 3 hpi). The phenomenon is consistent with others although the exact time it takes for upd3 to be induced varies depending on the tissues or the sensitivity of techniques employed (Pastor-Pareja *et al.* 2008; Ramond *et al.* 2020). The small delay in Upd3 expression compared with Toll/JNK activation, fits well with our model that *upd3* is transcriptionally controlled by Toll/Dl, and it then functions as a secondary cytokine signal. Importantly, injury-induced and Toll-dependent *upd3* expression in *Drosophila* hemocytes is very similar to the upregulation of secondary proinflammatory cytokine genes, such as IL-6 and IL-10, by TLR signaling in mammalian macrophages (O’Neill *et al.* 2013).

Collectively, injury-induced Toll, JNK, and JAK/STAT signaling in hemocytes leads to a number of intrinsic as well as systemic inflammatory responses. Within the hematopoietic system, one clear outcome is the differentiation of lamellocytes, and this work expands on early evidence (Márkus *et al.* 2005) to establish that the injury stimulus is sufficient to initiate and mediate lamellocyte formation even in the absence of wasp parasitization. Furthermore, activating mutations in the Toll and JAK/STAT signaling pathways (using *Tl^10b^* and *hop^Tum-l^*) elicit lamellocyte differentiation (Gerttula *et al.* 1988; Luo *et al.* 1995; Harrison *et al.* 1995; Govind 1996; Qiu *et al.* 1998). Our work shows that under normal genetic backgrounds, injury activates both Toll and JAK/STAT pathways and leads to lamellocyte differentiation.

The pathways discussed in this paper are broadly conserved (Matsuguchi *et al.* 2003; Pastor-Pareja *et al.* 2008; Jiang *et al.* 2009; Buchon *et al.* 2009b; reviewed in Kawasaki and Kawai 2014; Wu *et al.* 2015; Chakrabarti *et al.* 2016; La Marca and Richardson 2020). In the proposed model (Figure 7), we are able to place the events following injury in a temporal sequence based on observed phenotypes. The propagation of the injury-induced calcium wave within the epidermis is essentially instantaneous (Razzell *et al.* 2013). The next series of events, from calcium-dependent Duox activation and the production of hydrogen peroxide, to activation of Toll pathway and JNK pathway signaling in blood cells, all occur within the first few minutes to hours. The complete phenotypic effects of these events are observable within 3 hpi. Transcription of *upd3* and its activation of JAK/STAT pathway, require some more time, but these are fully apparent by 6 hpi. The consequences of these signals in mounting a cellular response is seen with full differentiation of lamellocytes within 24 hrs of the injury. In addition to the series of linked activation events (Figure 7), each individual pathway has its own downstream function in this inflammatory response. Mmp1, downstream of JNK, remodels the tissue surrounding the injury site (Stevens and Page-McCaw 2012; Purice *et al.* 2017). AMPs downstream of Toll anticipate any bacterial challenge (reviewed in Lemaitre and Hoffman 2007), and NFκB has multiple functions in inflammatory response (reviewed in Liu *et al.* 2017). In addition to lamellocyte formation, the Upd3 cytokine signal likely primes other tissues such as circulating hemocytes and fatbody for a possible innate immune response (Agaisse *et al.* 2003). Differentiation of *Drosophila* lamellocytes from macrophage-like cells (*i.e.* plasmatocytes) or progenitors is reminiscent of the differentiation of specialized macrophage classes in mammals in response to cytokine signaling. An example of this is the transformation of activated macrophages into non-phagocytic epithelioid cells during granuloma formation (Cronan *et al.* 2016). In the future, it will be important to determine if pro-inflammatory signaling in *Drosophila* involves prostaglandins and other eicosanoids, which have important roles in mammals (Rouzer and Marnett 2009; Medzhitov 2010). The role of inflammatory prostaglandin signaling has not yet been investigated in *Drosophila*, although a COX-like enzyme called Pxt has been identified (Tootle and Spradling 2008). Evolutionarily, a response to any breach of the body cavity precedes innate and acquired immune responses. Understanding the molecular events and their sequential function, we hope, will further mammalian studies on injury and healing.

**Figure 7.**
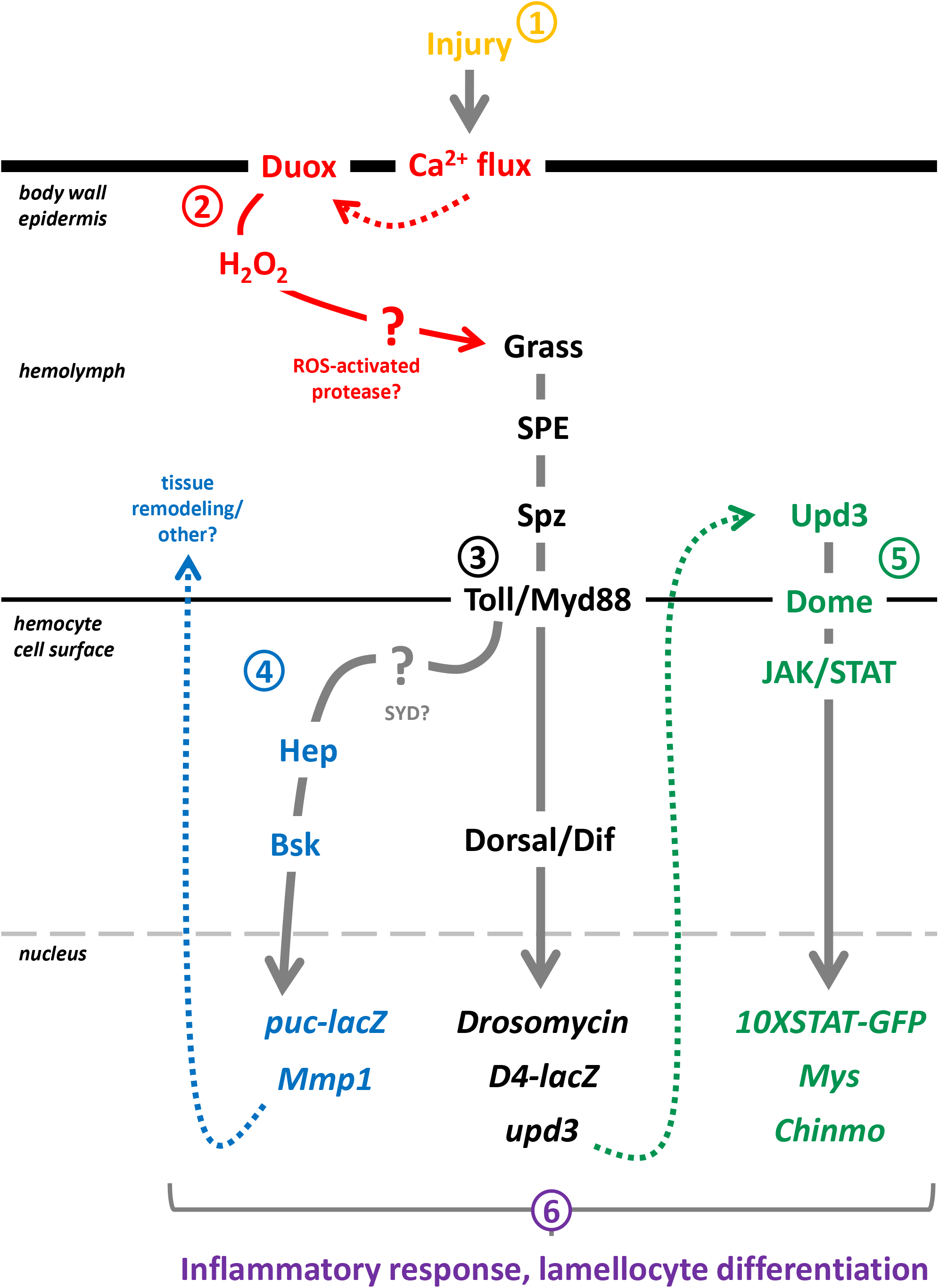
Model of injury-induced inflammatory signaling in the larval blood system. Injury to the larval epidermis (**1,** in yellow) leads to a wave of calcium signaling (Razzel *et al.* 2013) that activates epidermal Duox and produces ROS (**2**, in red) that activates the serine proteases Grass and SPE, which in turn, activate Spz and initiate Toll signaling in blood cells (**3**, in black). The Toll/Myd88 complex activates Dorsal, translocating it to the nucleus. Dorsal/Dif transcriptionally upregulates several Toll targets, including the cytokine-like *upd3* gene. Crosstalk between Toll and JNK signaling components JNK (Bsk) and JNKK (Hep) (**4**, in blue) at the level of the Toll receptor complex is proposed here based on published data on the scaffolding protein JIP3 (SYD) (Matsuguchi *et al.* 2003). As a result, the expression of JNK targets in the cell, including *puc* and *Mmp1* is Toll/Myd88 dependent. Mmp1 is a secreted protein that accumulates in circulating and lymph gland blood cells and upon secretion, functions in tissue remodeling, as cells are recruited to the injury site. Spz has been proposed to be a downstream target of JNK (Wu *et al.* 2015) allowing the possibility of a positive feedback loop in Toll activation following injury. After a short delay that allows synthesis and secretion of Upd3, this cytokine signals *via* its receptor, Dome and activates JAK/STAT target genes, including *mys* and *chinmo* (**5**, in green). Collectively, rapid Toll and JNK signaling, along with secondary JAK/STAT signaling, mediate the pro-inflammatory response to injury within the blood system and drive the differentiation of lamellocytes (**6**, in purple).

## Acknowledgments

We are grateful for the support provided to U.B. by NIH grants R01 HL-067395, R01 CA-217608, and R21 AI-094048; T.L. by a China Scholarship Council award; J.R.G. by Kirschstein National Research Service Award number T32 HL-69766 and UPLIFT (UCLA Postdocs’ Longitudinal Investment in Faculty) Award number K12 GM-106996. We thank Amy McWhorter for assistance with early injury experiments and discussion. We thank the members of the Banerjee laboratory, and in particular, acknowledge the contributions of Vivien Ho. We thank A. Courey, B. Lemaitre, J.M. Reichhart, Y.T. Ip, A. Page-McCaw, N. Perrimon, D. Stein, E. LeMosy, and S. Wasserman for fly stocks and advice. C.J.E. and U.B. devised the project. C.J.E., and T.L. performed experiments. C.J.E., J.R.G., and U.B. analyzed and interpreted data. C.J.E., J.R.G., and U.B. wrote the manuscript.

## Materials and Methods

### Fly Stocks

The following fly stocks were used in this work: *D4-lacZ* (A. Courey), *Drs-GFP, modSP^1^, Grass^Herrade^, SPE^Pasteur^, GNBP1^osiris^, GNBP3^Δ40^, psh^1^, spz^rm7^, da-gal4, Dif^2^* (B. Lemaitre), *Myd88^kra1^* (S. Wasserman), *Df(2L)J4, Df(2L)TW119* (Y.T. Ip), *ea^4^, snk^2^, gd^1^, ndl^10^* (D. Stein and E. Lemosy), *UAS-Duox-IR, UAS-Catalase, UAS-dome^DN^, 10xSTAT-GFP* (E. Bach), *upd3>GFP* (N.Perrimon), *Mmp1-lacZ* (D. Bohmann), *A58-gal4* (M. Galko), *Hml^Δ^-GAL4* (S. Sinenko), *UAS-Pxn, Toll^10b^*, *puc-lacZ* (U. Banerjee), αPS4-GFP (VDRC v318086), *Df(3L)ED4743 (GNBP1;* DrosDel), *Df(3L)ED4413 (GNBP3;* DrosDel), and each of the following from BDSC: *w^1118^*(stock 5905), *UAS-bsk^DN^, hep^r75^, Df(3R)Tl-P (spz, grass), Df(3R)Exel6205 (spz), Df(3R)Exel6195 (SPE), Df(3R)Exel6208 (grass), Df(3R)Exel6270 (modSP), Df(3R)BSC741* (*ea*), *Df(3R)Exel8157 (snk), Df(1)BSC542 (gd), Df(3L)BSC411 (ndl).*

### Axenic cultures and injuries

The axenic culture protocol was adapted from Brummel *et al.* (2004). Briefly, a sterile working environment was created by first washing a work hood with 70% ethanol followed by ultraviolet irradiation. Standard fly food was autoclaved and then poured into sterile vials or petri dishes inside the sterilized hood and allowed to cool before use. *Drosophila* embryos were collected from plates and washed with purified water using standard procedures, and then transferred to a 1.5 mL microfuge tube. Embryos were sterilized using two-fold diluted bleach followed by two washes in 70% ethanol and two washes in sterile water. Sterilized embryos were then transferred via sterile pipet to the previously prepared sterile food cultures. Fly cultures were grown at room temperature inside the sterilized work hood until the point of injury, and injured, sterile larvae were returned to the hood for recovery. For injury of axenic larvae, dissection plates were washed in 70% ethanol and UV irradiated, while forceps, pins, and pin holders were sterilized by autoclave. Water and 1X PBS were sterilized by vacuum filtration into sterile bottles. Cultures were verified as sterile using standard colony forming unit (CFU) assays sampling both cultures and larvae. Briefly, for cultures, 2 mL of sterile water was washed over the surface of the food for approximately one minute, then 1 mL was retrieved via sterile pipet to a sterile 1.5 mL microfuge tube. The microfuge tube was centrifuged briefly to pellet food debris, and 100 μL of supernatant was spread onto LB plates. For larvae, five larvae were collected in sterile water in a 1.5 mL microfuge tube and pulverized using a sterile, disposable micropestle. Larval carcass debris was pelleted by quick centrifugation and 100 μL of supernatant was spread onto LB plates. Standard fly cultures of similar developmental age were used as a positive control, while the sterile water vehicle alone served as a negative control. Seeded LB plates were sealed with Parafilm and left at room temperature for 4-5 days, by which point microbial colonies could be readily observed on positive control plates.

Wandering third instar larvae (for 3 hpi experiments) or early third instar larvae (for 24 hpi experiments) were removed from vials and washed thoroughly with purified water. Larvae were then transferred to a drop of 1X PBS pH 7.4 on a silicone dissection plate (Silgard). Individual larvae were gently stabilized dorsal-side up using forceps while a single puncture injury was carefully made to the lateral body wall at approximately 75% body length from the anterior. Puncture injuries were made using a sharp minutien pin (Fine Science Tools 26002-15) held in a pin holder (Fine Science Tools 26018-17). Immediately after injury, larvae were transferred to a standard food plate at room temperature for recovery.

### Dissections, bleeds, immunofluorescence, and cell counts

The dissection of larval lymph glands, the collection of circulating blood cells, and their analysis by fluorescence or immunofluorescence was performed using standard procedures, as previously described (Evans *et al.* 2014). For lamellocyte counting in lymph glands, dissected samples were either from the αPS4-GFP reporter line (Figure 3) or were immunostained for Atilla/L1 expression (Figure 6), then imaged via fluorescent confocal microscopy. Subsequent Z-stack image files were analyzed using ImageJ and the “3D” plug-in in order to more clearly visualize individual lamellocytes. Circulating cells were immunostained for Mys expression (Figure 3) on glass slides with circular “wells” created by a hydrophobic coating, then imaged using fluorescent confocal microscopy. The 20X objective was focused on the center of each circular well, and the field of view was captured as an image. The number of lamellocytes was counted for each image and used for statistical analysis. For immunostaining, the following antibodies were used: mouse anti-beta-galactosidase (Promega; 1:100), mouse anti-Mmp1 (1:1:1 mixture of DSHB 3A6B4, 3B8D12, and 5H7B11; 1:100), rabbit anti-Chinmo (E. Bach; 1:250), mouse anti-Mys (DSHB CF.6G11; 1:10), mouse anti-L1 (I. Ando; 1:100), mouse anti-Dorsal (DSHB 7A4; 1:10), mouse anti-Dif (Y. Engstrom; 1:250).

### Quantitative real-time PCR analysis

Circulating cells from ten larvae were isolated in 20 μL of 1X PBS for each biological replicate and total RNA was extracted using the PureLink™ RNA mini kit (Ambion) and quantitated using a spectrophotometer (Implen). The SuperScript® III First-Stand synthesis SuperMix kit (Invitrogen) and 150 ng of RNA was used for cDNA synthesis and relative quantitative PCR was performed by comparative C_T_ method using Power SYBR® Green PCR master mix kit (Applied Biosystems) with a StepOne™ Real-Time PCR detection thermal cycler (Applied Biosystems). Primers used in this study were either from published literature or designed using Primer3, and the expression level of *RpL32* was used to normalize total cDNA input in each experiment. Primer sequences (5’-3’) are as follows: *Drs*: (forward) CGTGAGAACCTTTTCCAATATGA, (reverse) TCCCAGGACCACCAGCAT; *Mmp1*: (forward) GGCAGAGGCGGGTAGATAG, (reverse) TTCAGTGTTCATAGTCGTAGGC; *upd*: (forward) AACTGGATCGACTATCGCAAC, (reverse) CTATGGCCGAGTCCTGGCTAC; *upd2*: (forward) CCAGCCAAGGACGAGTTATC, (reverse) GCTGCAGATTGCCGTACTC; *upd3*: (forward) ACAAGTGGCGATTCTATAAGG, (reverse) ATGTTGCGCATGTACGTGAAG; *Rpl32*: (forward) GACGCTTCAAGGGACAGTATCT, (reverse) AAACGCGGTTCTGCATGAG.

## Supplementary Figures

**Figure S1.**
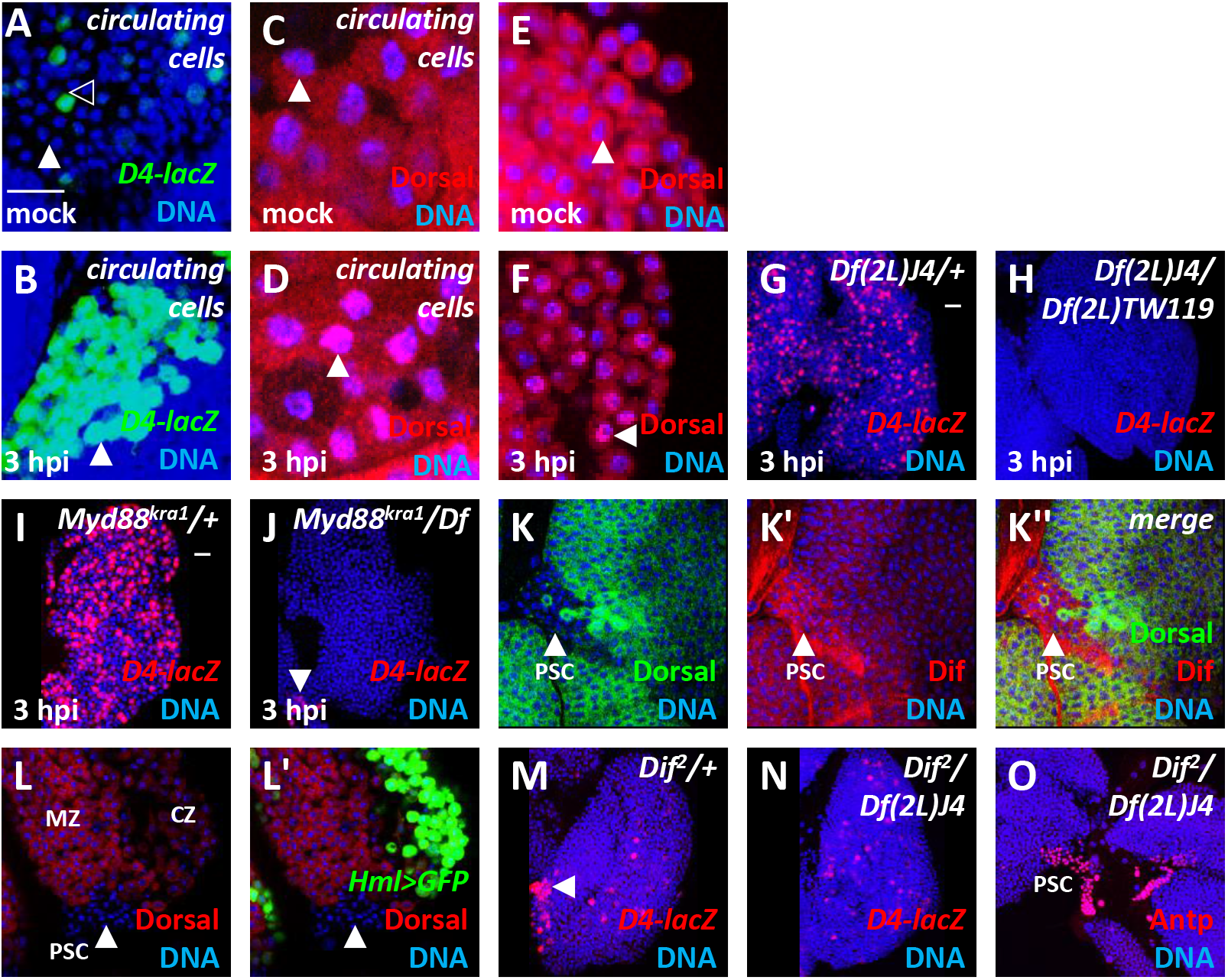
Distal epidermal injury activates Dorsal/Dif in the blood system. **(A-B)** *D4-lacZ* (green) and DNA (blue). Scale bars: 20 μm. **(A)** In mock (uninjured) larvae, very few circulating blood cells (shown are sessile eye disc cells) exhibit expression of *D4-lacZ* (black arrowhead). The white arrowhead points to a *D4-lacZ* negative cell. **(B)** At 3 hpi, circulating cells exhibit robust expression of *D4-lacZ* (white arrowhead). **(C-F)** Immunostaining of the NFκB-like transcription factor Dorsal (red); DNA (blue) in circulating **(C-D)** and lymph gland cells **(E-F).** Dorsal is present in the cytoplasm in mock (uninjured) larvae **(C, D)**, but by 3 hpi, blood cells accumulate Dorsal in the nucleus (white arrowheads). **(G-H)** Injury-induced *D4-lacZ* expression (red) in lymph gland cells 3 hpi in *dl/dif* double heterozygotes; the small J4 deletion removes both genes (**G)**, but *D4-lacZ* expression at 3 hpi is completely lost in homozygous Dorsal/Dif double null mutants (**H**; the *TW119* deletion overlaps with and larger than J4). Scale bars: 20 μm. **(I-J)** Loss of function in *Myd88 (Myd88^kra1^*) also suppresses injury-induced *D4-lacZ* expression (red), indicating its control of Dorsal/Dif activity. PSC cells continue to maintain *D4-lacZ* (white arrowhead in **J**). Scale bars: 20 μm. **(K-K’’)** Dorsal protein (**K, K’’**; green) is expressed in lymph gland primary and secondary lobe blood cells, but is absent from the PSC (arrowhead). Dif protein (**K’, K”**; red) is also expressed in lymph gland primary and secondary lobe blood cells but, in contrast to Dorsal, is also expressed in PSC cells at a relatively high level (arrowhead). **(L-L’)** Within the primary lymph gland lobe, Dorsal (red) is most highly expressed in the medullary zone (MZ) progenitor cells, with lower levels present in mature and maturing cortical zone (CZ) cells expressing the maturation marker Hemolectin (green, *Hml^Δ^-GAL4 UAS-GFP).* As in **(K)**, Dorsal protein is not observed in PSC cells (arrowhead). **(M-O)** Loss of *Dif* function **(N-O)** suppresses developmental *D4-lacZ* expression (red) in the PSC (arrowhead in **M**, missing in **N**) even though Dorsal-dependent expression is seen in the hematopoietic cells of the lymph gland. Immunostaining for Antp, a PSC marker, in *Dif* mutants (**O**, red) indicates that PSC cells are present in the *Dif* mutant backgrounds. In summary, Dif is exclusively expressed by and functions within PSC cells, while Dorsal and Dif both function in the blood cells. DNA is counterstained (blue) in all panels.

**Figure S2.**
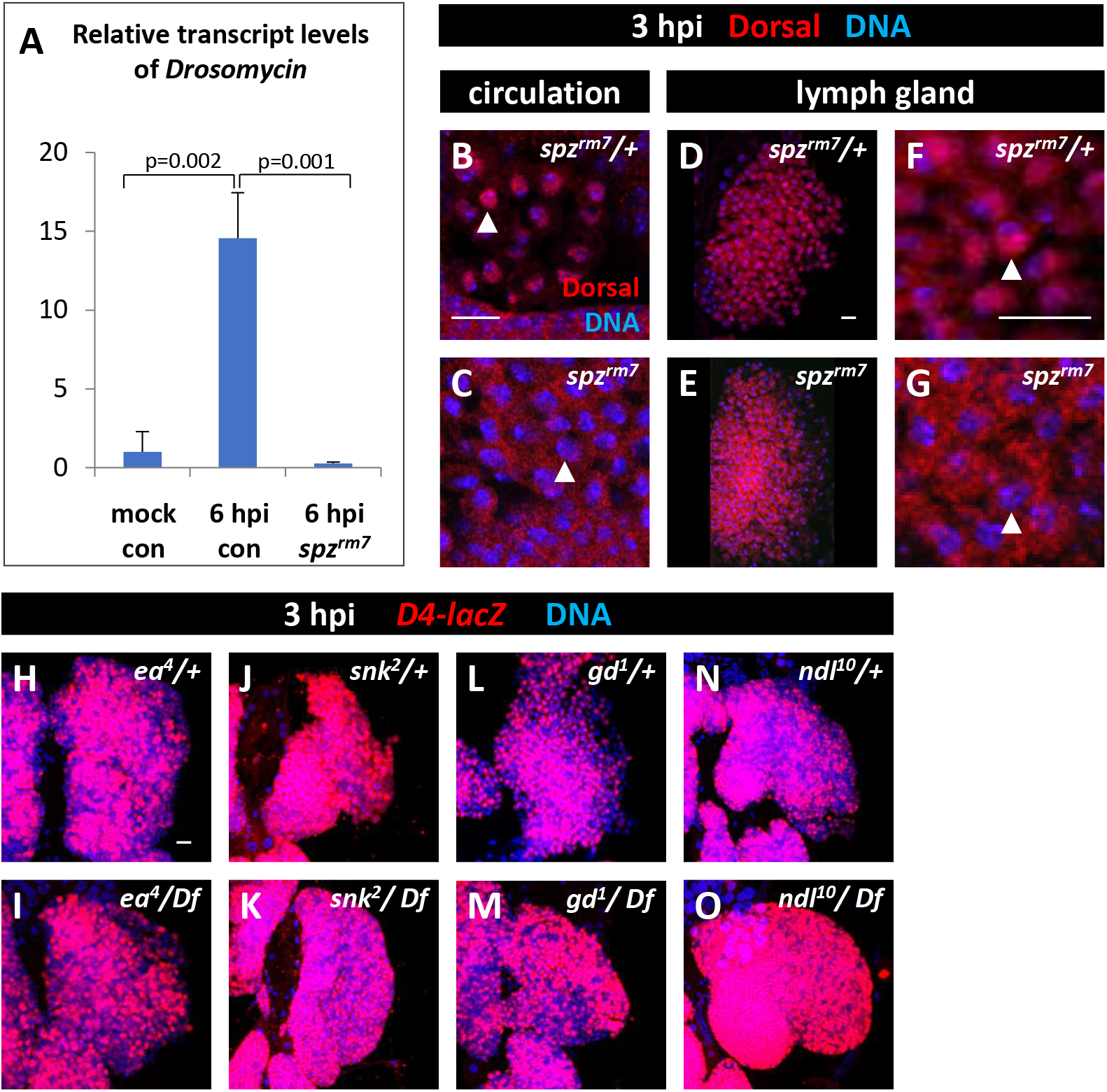
Injury-induced nuclear localization of Dorsal and expression of Drosomycin are dependent upon the Toll ligand Spz. **(A)** qRT-PCR analysis demonstrates that *Drosomycin* is upregulated by sterile injury approximately 15-fold by 6 hpi, and that the increase is suppressed in *spz^rm7^/spz^rm7^* mutants. **(B-G)** Immunostaining of Dorsal (red) at 3 hpi; DNA (blue); arrowheads indicate single nuclei. Genotypes are as indicated. Circulating cells **(B-C)** and lymph glands **(D-G)**; with higher magnification view in **(F-G)**. Scale bars 20 μm. Dorsal is localized to the nucleus of blood cells in injured heterozygous controls **(B, D, F)** but remains in the cytoplasm in injured homozygous *spz^rm7^* mutants **(C, E, G)**. **(H-O)** Loss of function of serine proteases that activate Spz during the specification of embryonic D/V polarity: *ea* (*easter*; **H-I**), *snk* (*snake*; **J-K**), *gd* (*gastrulation defective*; **L-M**), and *ndl* (*nudel*; **N-O**). None of these proteases suppress injury-induced *D4-lacZ* expression (red) in lymph glands at 3 hpi; DNA (blue). Scale bars: 20 μm.

**Figure S3.**
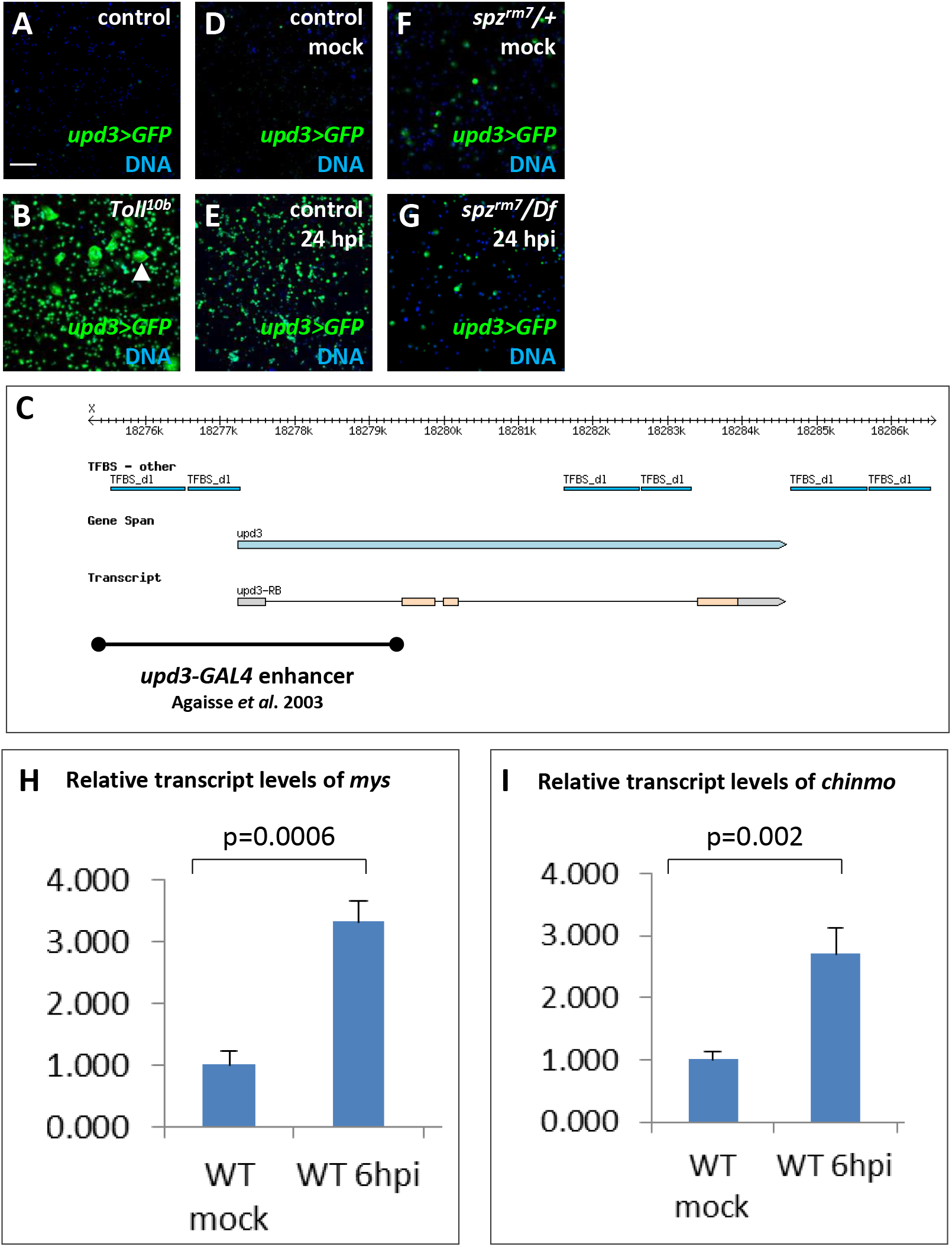
Regulation of *upd3* expression by Toll signaling. **(A-G)** *upd3* expression reported by *upd3-GAL4, UAS-GFP* (green; except in **C**). DNA (blue). **(A-B)** *upd3* is sparsely expressed by circulating cells in wild-type control **(A)**, but is strongly upregulated in *Toll^10b^* (*Tl^10b^*) gain-of-function mutants **(B)**. Note the presence of larger, flat lamellocytes in *Tl^10b^* mutants (arrowhead). Scale bars 100 μm. **(C)** The *upd3* locus (Flybase-GBrowser) displays transcription factor binding site (TFBS) regions for Dorsal (TFBS_dl). The black bar at the bottom of the schematic depicts the reported upd3 enhancer region used by Agaisse *et al.* (2003) in the construction of *upd3-GAL4.* **(D-G)** *upd3* is lowly expressed in circulating cells from uninjured larvae **(D)**, but sterile injury causes a large increase by 24 hpi **(E)**. This injury-induced increase is strongly suppressed by *spz^rm7^/Df* **(F-G)**. **(H-I)** qRT-PCR results showing that sterile injury induces expression of the JAK/STAT targets *mys* **(H)** and *chinmo* **(I)** at 6 hpi.

## Notes

### Competing Interest Statement

The authors have declared no competing interest.

